# Hydrogel-polyurethane fiber composites with enhanced microarchitectural control for heart valve replacement

**DOI:** 10.1101/2023.09.29.560202

**Authors:** Andrew Robinson, Abbey Nkansah, Sanchita Bhat, Shweta Karnik, Sarah Jones, Ashauntee Fairley, Jonathan Leung, Megan Wancura, Michael Sacks, Lakshmi Dasi, Elizabeth Cosgriff-Hernandez

## Abstract

Polymeric heart valves offer the potential to overcome the limited durability of tissue based bioprosthetic valves and the need for anticoagulant therapy of mechanical valve replacement options. However, developing a single-phase material with requisite biological properties and target mechanical properties remains a challenge. In this study, a composite heart valve material was developed where an electrospun mesh provides tunable mechanical properties and a hydrogel coating confers an antifouling surface for thromboresistance. Key biological responses were evaluated in comparison to glutaraldehyde-fixed pericardium. Platelet and bacterial attachment were reduced by 38% and 98%, respectively, as compared to pericardium that demonstrated the antifouling nature of the hydrogel coating. There was also a notable reduction (59%) in the calcification of the composite material as compared to pericardium. A custom 3D printed hydrogel coating setup was developed to make valve composites for device-level hemodynamic testing. Regurgitation fraction (9.6 ± 1.8%) and effective orifice area (1.52 ± 0.34 cm^2^) met ISO 5840-2:2021 requirements. Additionally, the mean pressure gradient was comparable to current clinical bioprosthetic heart valves demonstrating preliminary efficacy. Although the hemodynamic properties are promising, it is anticipated that the random microarchitecture will result in suboptimal strain fields and peak stresses that may accelerate leaflet fatigue and degeneration. Previous computational work has demonstrated that bioinspired fiber microarchitectures can improve strain homogeneity of valve materials toward improving durability. To this end, we developed advanced electrospinning methodologies to achieve polyurethane fiber microarchitectures that mimic or exceed the physiological ranges of alignment, tortuosity, and curvilinearity present in the native valve. Control of fiber alignment from a random fiber orientation at a normalized orientation index (NOI) 14.2 ± 6.9% to highly aligned fibers at a NOI of 85.1 ± 1.4%. was achieved through increasing mandrel rotational velocity. Fiber tortuosity and curvilinearity in the range of native valve features were introduced through a post-spinning annealing process and fiber collection on a conical mandrel geometry, respectively. Overall, these studies demonstrate the potential of hydrogel-polyurethane fiber composite as a heart valve material. Future studies will utilize the developed advanced electrospinning methodologies in combination with model-directed fabrication toward optimizing durability as a function of fiber microarchitecture.

## 1. Introduction

Over 290,000 patients require heart valve replacements worldwide, with an estimated increase to 850,000 by 2050.^1,2^ Current replacement heart valve options include mechanical valves and tissue-based bioprosthetic valves. Although mechanical valves provide excellent durability, patients are required to be on lifelong anticoagulants.^3–5^ Bioprosthetic valves are often preferred due to favorable hemodynamics and no long-term requirement of anticoagulant therapy.^6,7^ Unfortunately, the bioprosthetic valve has limited durability with 50% failing within 15 years due to structural degeneration resulting from material fatigue and calcification.^2,7,8^ Researchers continue to pursue polymeric valves as an alternative clinical option that offer the potential to combine the durability of mechanical valves and the hemocompatibility of bioprosthetic valves.^6^

Polymeric valves have been investigated since the early 1960’s offering a high degree of design freedom, reduced valve-to-valve variability, and reduced cost compared to bioprosthetic valves.^9,10^ In design of a polymeric valve, a biomaterial is required that can provide near physiological hemodynamics, mitigates thrombosis and calcification, and is engineered to resist material fatigue and degradation. Polymers under investigation have included silicone, polytetrafluoroethylene (PTFE), polyvinyl alcohol (PVA), and polyurethanes. Thrombogenicity of silicone valves resulted in early failure due to embolic events.^11,12^ Although versions of PTFE have been successful for use in vascular grafts, its use as a heart valve resulted in failures due to calcification and leaflet fragmentation.^13,14^ PVA valves have reported promising mechanical properties; however, their functional assessments and long-term durability require further investigation.^15^ Early polyurethane valves, such as poly(ester urethanes) and poly(ether urethanes), suffered from degradation-based failure.^6^ Although modern chemistries have improved the biostability of polymers, it remains challenging to achieve the requisite durability while limiting thrombosis and calcification in a single-phase material.

To decouple the biological and durability design goals, we have previously developed composite materials composed of a hydrogel-coated electrospun mesh wherein the mesh provides bulk mechanical properties and the hydrogel provides an antifouling surface.^16^ Compared to common industrial fabrication techniques such as molding, electrospinning produces non-woven fibrous meshes with a high degree of tunability over fiber architecture to control mechanical properties.^17–19^ Electrospun polyurethanes have demonstrated degradation resistance as well as elastomeric and bulk mechanical properties similar to cardiovascular tissues.^16,20^ Our prior evaluation of a randomly-oriented electrospun poly(carbonate urethane) fiber mesh, demonstrated reproducible mechanical behaviors beyond the estimated maximum physiological heart valve stresses highlighting its potential as a heart valve leaflet material.^21^ Coating of the polyurethane mesh with a PEG-based hydrogel provided an antifouling surface that resisted thrombus as compared to clinical materials such as ePTFE.^16,22–24^ The benefit to resisting thrombus extends to calcification that has been correlated with surface microthrombi in polyurethane materials.^25,26^ Although surface properties were modified utilizing the coating, mechanical evaluation of composites demonstrated that the hydrogel negligibly contributed to the bulk mechanical properties.^21^ This key finding indicates that we can independently optimize each phase without detriment to the other design goals. Thus, efforts to improve the thromboresistance and bioactivity of the hydrogel can proceed in parallel to efforts to improve durability of the electrospun mesh.

In contrast to the isotropic behavior of random electrospun meshes, native heart valve leaflets are highly anisotropic and have a non-linear mechanical response. From a structural framework, valve microarchitecture include fibers that are highly aligned, have a change in the main angle of alignment over the width of the valve from commissure to commissure (curvilinear), and have a crimped or tortuous morphology.^27,28^ Computational simulations indicate that reproducing these native fiber features improves strain uniformity with a hypothesized correlation to enhanced valve durability.^29^ Investigators have studied reproducing these native features with electrospun fiber microarchitecture and the resulting effect on heart valve performance.^21,30,31^ For example, fiber alignment imparts anisotropic mechanical properties more similar to native valve stress responses.^21,22,32^ Similarly, Hobson et al. used a conical collector to achieve fiber curvilinearity similar to the native heart valve with associated improvements in strain homogeneity at the device level compared to linearly aligned fibers.^28^ However, simple recreation of valve-inspired microarchitectures may not produce optimal durability of a synthetic valve due to the inherent differences between the synthetic polymer and extracellular matrix properties. Thus, to realize the potential of this approach, new methodologies are required to expand the level of fiber microarchitecture control to match or exceed physiological values of alignment, curvilinearity, and tortuosity observed in native tissue.

The current study first examines the biological and device-level hemodynamic performance of a hydrogel-polyurethane fiber composite valve. To confirm the antifouling surface properties, bacterial and platelet attachment to the composite specimens were tested *in vitro.* Composite calcification propensity compared to glutaraldehyde-fixed pericardium was also characterized. A custom 3D printed setup consisting of a chamber and hanger was then developed to apply unform coatings to device-level electrospun tubes using our redox-based coating platform.^33^ The hydrogel-polyurethane fiber composite tubes were sutured to a cobalt chromium stent for device level testing of hemodynamics. To enhance long-term durability, electrospinning methodology was developed to improve control of fiber alignment, curvilinear alignment, and tortuous microarchitectures. Collectively, this work provides preliminary assessment of a hydrogel-polyurethane fiber composite for use as a polymeric heart valve and the development electrospinning methodologies to create bioinspired microarchitectures for improve durability.

## 2. Materials and Methods

### 2.1 Materials

All chemicals were purchased from Sigma Aldrich (Milwaukee, WI) or VWR (Radnor, PA) and used as received unless stated otherwise.

### 2.2 Biological Responses of Composite Heart Valve Material

Calcification, platelet attachment, and bacterial adhesion of the hydrogel-polyurethane fiber composite valve material were evaluated and benchmarked against glutaraldehyde-fixed pericardium (clinical control). Fresh bovine pericardium was received from Animal Technologies (Tyler, TX) in saline and sectioned into approximately 2 cm x 2 cm squares. The sections were fixed in 0.5% glutaraldehyde for 72 hours at a concentration of 30 ml/g.^34,35^ Following the 72 hour fixation, samples were transferred to a 0.2% glutaraldehyde solution for one week and stored in the solution until use.^35,36^

#### 2.2.1 Flat Electrospun Mesh Fabrication

For initial benchtop testing of composite materials, meshes were electrospun from Bionate® 80A (DSM Biomedical Inc, Berkeley, CA, USA) that was solubilized in 70:30 dimethylacetamide:tetrahydrofuran. The electrospinning solution was pumped at a rate of 0.5 ml/hr through a 20-gauge blunt needle that was charged to +14.0-19.2 kV. Fibers were collected on a 26 mm diameter mandrel (40 mm in length) rotating at a velocity of 50 RPM and charged to –3.5-5.0 kV producing a random fiber orientiation. A working distance of 50 cm was utilized. Fibers were collected for 3 - 4 hours. Relative humidity in the spinning chamber was controlled between 44-50% with a temperature range of 19-24 °C. Upon completion of the spin, meshes were annealed at 70 °C overnight prior to removal from the mandrel. Meshes were anaylzed via scanning electron microscopy (SEM, Phenom Pro, NanoScience Instruments, Phoenix, AZ) at an accelerating voltage of 10 kV. Prior to imaging, samples were mounted and coated with 4 nm of gold (Sputter Coater 108, Cressington Scientific Instruments, Hertfordshire, UK).

#### 2.2.2 Flat Composite Fabrication

Polyether urethane diacrylamide (PEUDAm) was synthetized as previously described.^33^ Diffusion-mediated redox crosslinking was performed to hydrogel coat the electrospun polyurethane meshes.^23,33^ Briefly, mesh substrates (0.18-0.24 mm thick) were cut to 1.2 x 1.2 cm squares or 2.2 x 2.2 cm squares and soaked in a wetting ladder of decreasing concentrations of ethanol (70 vol%, 50 vol%, 30 vol%, and 0 vol% ethanol in water, 15 minutes each). Iron gluconate was then adsorped onto the substrates by immersing the meshes in iron gluconate dihydrate solutions (IG, 3 wt/vol% [Fe^2+^], as determined with the Ferrozine Assay^37^) for 15 minutes. Substrates were briefly dipped in methanol (< 2 seconds) and dried for one minute under compressed air. Meshes were then placed in 3D printed clamps to prevent folding and submerged in a polymer precursor solution comprised of 10 wt/vol% 20 kDa PEUDAm and 0.05 wt/vol% ammonium persulfate (APS) for 20 seconds. Hydrogel-coated composites were air dried overnight then swelled in DI water for 5 hours with water exchanges at 10 minutes, 1 hour, and 3 hours to remove unincorporated polymer and residual redox species. For improved toughness, a second interpenetrating network was incorporated based on our previous report.^33^ The dried composites were swollen in a solution of 20 wt/vol% N-acryloyl glycinamide (NAGA; BLD Pharma), 0.1 mol% bisacrylamide (bisAAm), and 2 wt/vol% Irgacure 2959 overnight protected from light at 4 °C. Swollen substrates were placed on a glass plate and cured for 20 minutes on each side via photopolymerization under a UV transilluminator (Intelligent Ray Shuttered UV Flood Light, Integrated Dispensing Solutions, Inc., 365 nm, 4 mW/cm^2^). Composites were then soaked in deionized water to remove unreacted monomer.

#### 2.2.3 Calcification

A 2x simulated body fluid (SBF) was prepared by adapting a protocol from Kokubo et al.^38^ Briefly, 700 ml of DI water was stirred at 37 °C with pH and temperature monitored using a Sartorius pHBasic pH meter. Sodium chloride (275.0 mM), sodium bicarbonate (8.5 mM), potassium chloride (6.0 mM), potassium phosphate dibasic anhydrous (2.0 mM), and magnesium chloride anhydrous (3.1 mM) were added in their respective order with each allowed to dissolve completely prior to the addition of the subsequent reagent. To facilitate calcium dissolution, 78 ml of hydrochloric acid (1.0 M) was added. Calcium chloride (5.3 mM) and sodium sulfate (1.1 mM) were then individually dissolved in the solution. Deionized water was added to bring the solution volume to 900 ml. The solution was then titrated to a pH of 7.20 ± 0.01 with tris hydroxymethyl aminomethane that had been dissolved in 100 ml of deionized water (1.0 M) and sterifiltered.

Prior to the calcification study, the fixed pericardium underwent four washes in deionized water for 30 minutes each and an overnight wash to remove excess fixative. Composites were swollen in deionized water overnight to ensure equilibrium. All samples were cut to 1 cm x 1 cm and vertically suspended in a 50 ml conical tube with monofilament and a Teflon weight. Samples were submerged in the 2x SBF with solution changes occurring every other day for two weeks. Following the two-week study, specimens underwent four washes in DI water for 30 minutes each to remove non-nucleated species. Specimens were randomly assigned to either quantification or morphological assessment under scanning electron microscopy. Pericardium specimens for SEM analysis underwent a serial dehydration in 30%, 50%, 70%, 90%, 95%, and 100% ethanol for 10 minutes each.^39^ Following the serial ethanol dehydration, the samples were dried in 50:50 ethanol:hexamethyldisilazane (HDMS) for 10 minutes and then in pure HDMS and allowed to air dry overnight.^39^ Hydrogel-polyurethane fiber composites for SEM were air dried overnight to mitigate curling. Samples were sputter coated with 4 nm of gold as described and analyzed on a Thermo Fischer Scientific Apreo 2 SEM at a 10 kV accelerating voltage. The remaining specimens were vacuum dried overnight for quantification. Vacuum dried samples were massed and hydrolyzed in 250 µl of 5.0 N HCl for six hours.^40^ Calcium content within the lysate was quantified using the o-cresolphthalein complexone (n = 5) according to manufacturing instructions (Calcium LiquiColor® Test, Stanbio Laboratory) with absorbance measured at 550 nm (Tecan Infinite Nano+). The calcium content was normalized to dry mass and the deposition of calcium was calculated by subtracting the calcium content of untreated specimens from the SBF treated specimens.

#### 2.2.4 Platelet Attachment

As an initial assessment of thromboresistance, static platelet attachment was characterized as previously described in literature.^33,41^ Glutaraldehyde-fixed pericardium and hydrogel-polyurethane fiber composites were prepared by immersion in 70% ethanol for 3 hours. Samples were then subjected to a wetting ladder with sequential soaks in 50%, 30% and 0% ethanol in PBS for 30 min each. Specimens (8 mm discs, n = 5) were then soaked in FBS overnight at 37 °C, washed once with PBS to remove non-adhered protein, and transferred to a new 48 well plate. Platelets were isolated from human whole blood drawn in a tube with heparin from a healthy volunteer from which informed written consent was obtained (IRB protocol STUDY00002311). Platelet rich plasma (PRP) was isolated by centrifuging the whole blood at 990 RPM for 15 minutes. The PRP layer was removed, and prostacyclin was added at 10 μL/mL. The PRP was re-centrifuged at 1500 RPM for 10 minutes. The obtained pellet was resuspended in CGS buffer and centrifuged at 1500 RPM for 10 minutes. Using Tyrode’s buffer, the platelet pellet was resuspended at half the original volume of PRP. Sudan B Black (5% in 70% ethanol) was added at a ratio of 1:10 to the platelet solution and incubated for 30 minutes at room temperature. The stained platelets were washed three times with PBS then centrifuged at 1500 RPM for 8 minutes. Platelets were counted via a hemocytometer and resuspended in PBS at a concentration of 10 x 10^6^ platelets/mL. Specimens were then incubated with 300 μL of the platelet suspension for 30 minutes at 37 °C in a shaking incubator. Samples were carefully transferred to a new well, washed with PBS twice, and then retransferred to a new plate to eliminate non-adhered platelets and platelets that may be adhered to the well plate. Samples were treated with 150 μL of DMSO for 15 minutes to lyse adhered platelets with 150 μL of PBS then being added. The absorbance of the lysed solution was read on a spectrophotometer (Tecan Infinite M Nano+) at 610 nm to quantify platelet attachment.

#### 2.2.5 Bacteria Adhesion

Bacterial resistance of the pericardium and hydrogel-polyurethane fiber composites were determined via characterization of static bacterial adhesion. In preparation for bacterial seeding, each sample was cut into 8 mm discs, sterilized by immersion in 70% ethanol for 3 hours, and subjected to a wetting ladder with sequential soaks in 50%, 30% and 0% ethanol in PBS for 30 minutes each. A non-specific strain of methicillin-resistant staphylococcus aureus (ATCC 25923) was utilized for the studies due to the increasing association of MRSA in bacterial endocarditis and the high morbidity rate estimated at 40-80%.^42^ The bacteria culture was expanded overnight, and specimens were incubated in an orbital incubator with 600 µl a bacterial suspension of 1 x 10^7^ CFU/mL for 2 hours at 37 °C and 120 rpm. Non-adhered bacteria were removed by rinsing in PBS. The remaining adhered bacteria were removed via alternating sonication for five minutes and vortexing for 10 seconds, 3 times. The supernatant was plated on Brain Heart Infusion agar and incubated at 37 °C for 16-18 hours. Resulting colony forming units (CFU) were counted (n = 9). For the pericardium samples, the supernatant was diluted 3-fold in PBS before plating to enable an adequate density for counting. In preparation for imaging, all samples were fixed in 3.7% glutaraldehyde for 10 minutes and washed in 0.15 M NaCl buffer. The samples were incubated with 25 µg/mL wheat germ agglutinin CF™594 conjugate (Biotium; Excitation/Emission: 593/614 nm) diluted with NaCl buffer for 30 minutes at room temperature and subsequently washed with the NaCl buffer. Specimens were imaged with a fluorescent microscope (Nikon Eclipse TE2000-S). Specimens were then washed in deionized water 3 times for 20 minutes each (120 rpm, 37°C) to remove salts and allowed to air dry overnight prior to imaging with SEM.

### 2.3 Heart Valve Fabrication and Hemodynamic Assessment

Device level testing on hydrogel-polyurethane fiber composites with a random fiber microarchitecture was performed to measure baseline hemodynamic properties.

#### 2.3.1 Fabrication of Electrospun Heart Valve Tubes

Heart valve tubes were electrospun from a 14 wt% Bionate® 80A (DSM Biomedical Inc, Berkeley, CA, USA) solution in 70:30 dimethylacetamide:tetrahydrofuran. The solution was pumped at a rate of 0.5 ml/hr through a 20-gauge blunted needle that was charged to + 13.5-14.9 kV. Meshes with a random fiber orientation were fabricated on a 26 mm diameter mandrel (40 mm in length) that was charged to −2.5 kV at a working distance of 50 cm and a rotational velocity of 50 RPM. Fibers were collected for 3.5 - 4 hours. Relative humidity in the spinning chamber was controlled between 42-46% with a temperature range of 23-24°C. Upon completion of the spin, the meshes were annealed while on the mandrel at 70 °C overnight. Meshes were removed from the mandrel without cutting to maintain a tube, **Figure 3A**. Sections were taken from the tubes and prepared for SEM as previously described. Fiber diameter was characterized by drawing a mid-line through the image and measuring the diameter of the first ten fibers that crossed the line (n = 150 per mesh, mesh = 3). Mesh thickness was measured at nine location along the tube using force-normalized calipers, Mitutoyo 547-500S.

#### 2.3.2 Suture Retention Strength of Heart Valve Mesh

Suture retention strength of the electrospun heart valve material was tested according to ISO 7198 straight across procedure using an Instron 3345.^43^ Briefly, specimens were cut to 4 mm x 10 mm rectangular sections from tube specimens. A 5-0 PROLENE suture was inserted through the material 4 mm from the top edge forming a half loop. The suture was pulled at a uniaxial strain rate of 100 mm/min until the suture pulled through the material.^44^ The suture retention strength was recorded as the maximum force during testing (n = 5).

#### 2.3.3 Hydrogel Coating of Heart Valve Composites

Prior to coating, electrospun tubes were ramped in ethanol (70 vol%, 50 vol%, 30 vol%, and 0 vol% ethanol in water, 15 minutes each) and then submerged in a 3 wt% IG solution for 15 minutes, **Figure 1A**. To facilitate thin, uniform coatings of the first hydrogel network, a custom 3D printed chamber and hanger system was designed, **Figure 1B**. A chamber with an internal diameter of 30 mm, a central pillar 18 mm in diameter, and a height of 40 mm was utilized to hold the PEUDAm and APS solution. The central pillar reduced the volume of precursor solution required. A hanger system consisting of two rings was designed to guide and aid in stabilization of the tube during coating. The dimensions of the ring were chosen so that they would fit inside the tube without shifting during handling, but not stretch the material. The bottom ring had an outer diameter of 24 mm, was 1.5 mm thick, and had a height of 2 mm. The upper ring had the same dimensions but had an upper attachment for handling. The rings were positioned inside the electrospun tube as shown in **Figure 1C**. The tube was then guided into the chamber using the fitted hanger system that held a solution of 10 wt/vol%, 20 kDa PEUDAm, 0.05 wt/vol% ammonium persulfate, and 2 wt/vol% Irgacure 2959 for 20 seconds. Irgacure 2959 was added to increase the degree of crosslinking. Immediately following the 20 second immersion, the composites were photopolymerized on a UV plate (Intelligent Ray Shuttered UV Flood Light, Integrated Dispensing Solutions, Inc., 365 nm, 4 mW/cm^2^) for 12 minutes and allowed to dry overnight. After drying, composites were soaked for 5 hours in DI water to remove the sol fraction. Water exchanges were performed after 10 minutes, 1 hour, and 3 hours. Composites were then dried again overnight. The second interpenetrating hydrogel network was introduced by soaking dried composites in a solution of 20 wt/vol% pNAGA, 0.1 mol% bisAAm, and 2 wt/vol% Irgacure 2959 overnight protected from light at 4 °C. Composites were then photopolymerized on a UV plate (Intelligent Ray Shuttered UV Flood Light, Integrated Dispensing Solutions, Inc., 365 nm, 4 mW/cm^2^) for 40 minutes and swollen in DI water, **Figure 1D**. After overnight swelling, hydrogel thickness was measured at the top, middle, and bottom of the composite three times in each region with Mitutoyo 547-500S force-normalized calipers to evaluate coating uniformity.

**Figure 1:**
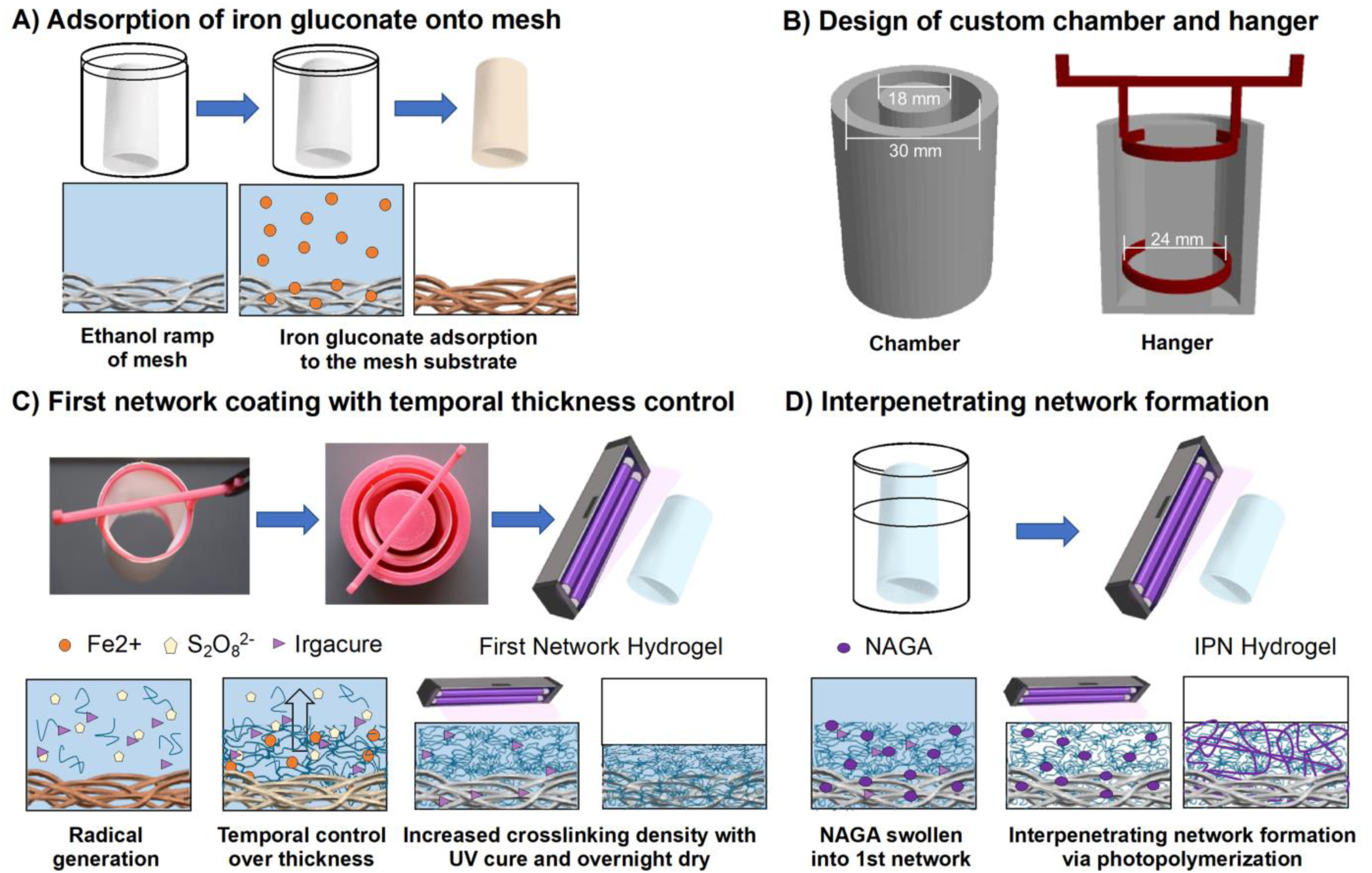
Schematic of 3D printed coating setup for the fabrication of hydrogel-polyurethane fiber composite valve tubes. A) Ethanol ramp for wetting and iron gluconate adsorption onto the electrospun mesh. B) 3D printed chamber and hanger system. The chamber with center pillar reduces the amount of polymer with the hanger providing stability during the coating process. C) Dipping of the iron gluconate adsorbed mesh in the hydrogel precursor solution. The hydrogel coating thickness increases with submersion time. A UV photocuring step was added to increase crosslinking of the first network. D) Swelling of the second network into the primary composite with UV photocuring resulting in final IPN composites.

#### 2.3.4 Heart Valve Assembly

A custom-made stent design was manufactured using a 26 mm cobalt-chromium (Co-Cr) alloy stent, a common medical grade alloy currently commercially used in FDA approved heart valve devices.^45–47^ The custom stent was placed inside the hydrogel-polyurethane fiber composite tube and commissural points were identified. The mid-points of the composite tube between the commissures were brought to the center manually to ensure that the material was properly coapting. Two sutures were placed on each commissure (six total) to hold the material in place thereby forming the three leaflets from the single tube material. A single running stitch was placed on the circumference of the stent frame, following the frame along the leaflet attachment points and curvature to secure the hydrogel-fiber composite tube forming the heart valve. The stent was crimped to ∼ 23 mm diameter to ensure the leaflet edges touched each other sufficiently at the inflow section to prevent experimental regurgitation but avoid leaflet pinwheeling. The valve was hydrated throughout the suturing process to prevent drying of the hydrogel.

#### 2.3.5 Hemodynamic Testing

The valve prototype was tested in a custom left heart simulator to evaluate the hemodynamic characteristics of the prototype (n = 3). The simulator replicates the pulsatile motion of the left heart and is seeded with a 60/40 mixture of water/glycerin (99% pure glycerin) to mimic blood viscosity. Further details on the simulator are described in previous studies. ^48–51^ The effective orifice area (EOA) is an important parameter to evaluate valve orifice opening in addition to the efficiency of the valve.^52^ EOA was computed using the Gorlin’s equation (Eq. 3):

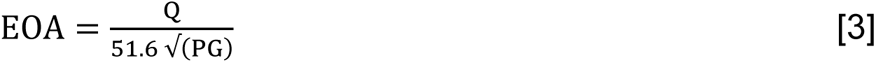

Where Q represents the root mean square aortic valve flow over the same averaging interval of the pressure gradient, ΔP. Regurgitant fractions (RF) were calculated as the ratio of the closing volume (CV) and leakage volumes (LV) to the forward flow volume (FV) as follows:

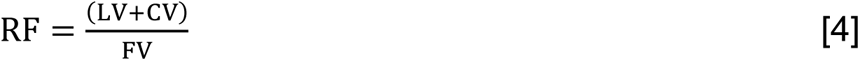

The mean pressure gradient was calculated as the difference between the upstream and downstream pressures across the valve. Data was calculated by averaging metrics over 100 consecutive cardiac cycles.

### 2.4 Microarchitectural Control

Electrospinning was utilized to develop methodologies to expand control of material microarchitecture to produce bioinspired ranges of fiber alignment, curvilinearity, and tortuosity.

#### 2.4.1 Alignment

An electrospinning solution of 14 wt% Bionate® 80A (DSM Biomedical Inc, Berkeley, CA, USA) in 70:30 dimethylacetamide:tetrahydrofuran was used for alignment investigations. The electrospinning solution was passed through a 20-gauge needle tip at 0.5 ml/hr flow rate with an applied positive voltage of 11.2-16.0 kV. A 4 cm diameter mandrel with an applied charged of −5 kV was utilized to collect fibers at a distance 50 cm from the charged needle tip. Humidity was controlled to 38-40% and temperature was ambient, ∼21 °C. Rotational velocities of 50 RPM and 4000 RPM were investigated. Samples were prepared and anaylzed via scanning electron microscopy as previously described. Fiber alignment was quantified utilizing a semi-automated algorithm written in Matlab (The Mathworks, Natick, MA) by D’amore et al. to determine the normalized orientation index (NOI) (n = 5).^53^ Values closer to zero indicate a random fiber orientation and those closer to 100 indicate a highly aligned orientation.

#### 2.4.2 Curvilinearity

Conical mandrels were investigated as a means to introduce curvilinearity to meshes based on the work by Hobson et al.^28^ An electrospinning solution of 14 wt% Bionate® 80A (DSM Biomedical Inc, Berkeley, CA, USA) in dimethylacetamide was utilized. The solution was passed through a charged (+15 kV) 20-gauge needle at a flow rate of 0.5 ml/hr. Humidity ranged from 36-48% and the temperature was 20-22 °C. A 3D printed conical mandrel was designed with a base diameter of 7.5 cm and a height of 3.5 cm. A cylindrical mandrel with a diameter of 4 cm was utilized as a control for linear aligned fibers. The mandrels were charged to −5 kV and rotated at 4000 RPM at a distance 35 cm from the needle tip. Mesh sections were taken from the middle of the cylindrical mandrel along the fiber alignment. Sections ∼15 mm wide by ∼10 mm high were taken 1.5 cm from the base of the conical mandrel with the width of the samples along the direction of alignment. Sectioning 1.5 cm from the bottom of the mandrel was chosen based on reported literature leaflet heights for native and bioprosthetic valves as a means to look at the change in the main angle of alignment across the top (free edge) of the theoretical leaflet.^54,55^ Samples were prepared for SEM as described above. Images were taken at 0, 5, 10 mm centered on the 1.5 cm strip and at the top of the sections (1.5 cm from the base of the mandrel) along the direction of alignment. The directionality plugin in ImageJ was utilized to determine the main angle of alignment. The degree change per mm was calculated by determining the difference between the images taken at 0 mm and at 10 mm and then dividing by 10 (Eq. 1) (n = 25).

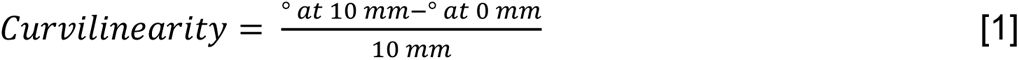

#### 2.4.3 Tortuousity

A solution of 14 wt% Bionate ® 80A (DSM Biomedical Inc, Berkeley, CA, USA) in dimethylacetamide was used for electrospinning. The solution was passed at a flow rate of 0.5ml/hr through a 20-gauge needle tip that was charged to 12.5 kV. A working distance of 35 cm was used. Humidty and temperature were ambient, 40% and 22-23 °C. Fibers were collected on a 4 cm diameter rotating mandrel at 4000 rpm. The mesh was split along the circumference of the mandrel with one of the pieces left on the mandrel and one piece removed from the mandrel. Both sheets were annealed at 70 °C overnight. Samples were prepared for SEM as previously described. Tortuosity was manually analyzed using Image J. A horizontal line was drawn across the image and the tortuosity of the first ten fibers that crossed the line were measured according to Eq. 2, below (n = 30).

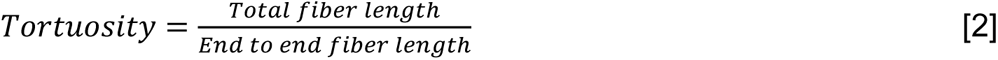

### 2.5 Statistical Analysis

Data are represented as mean ± standard deviation, unless otherwise stated. Statistical analysis was performed using a t-test analysis in Prism GraphPad unless stated otherwise. Statistical significance was set at p < 0.05.

## 3. Results

### 3.1 Biological Responses to Heart Valve Composite

Biological responses of the hydrogel-polyurethane fiber composite were evaluated in comparison to glutaraldehyde-fixed pericardium that served as a clinical benchmark. Calcification propensity was evaluated using a 2x simulated body fluid, **Figure 2A**. Morphological assessments via SEM displayed deposits on the treated pericardium specimens, whereas composite specimens displayed minimal deposition similar to non-treated controls, **Figure S1**. The o-cresolphthalein assay was used to quantify the calcium deposition on the SBF treated samples and normalized to non-treated specimens. The flat composite (2.5 ± 0.3 µg/mg sample) had a marked reduction in calcium deposition as compared to the pericardium specimens (6.1 ± 0.7 µg/mg sample) (59% reduction, *p* < 0.00001, *n* = 5). Bacterial and platelet adhesion was utilized to evaluate the antifouling nature of the hydrogel coating. Hydrogel-polyurethane fiber composites demonstrated a significant reduction in platelet attachment (2.6 x 10^6^ platelets/cm^2^) as compared to the pericardium (4.3 x 10^6^ platelets/ cm^2^) (38% reduction, *p* < 0.0001, *n* = 5), **Figure 2B**. Similarly, a reduction in bacteria for the composite was observed in fluorescent and SEM images, **Figure 2C**. Quantification of the colony forming units demonstrated a significant reduction in attached bacterial CFUs for the flat composite specimens as compared to the pericardium (98% reduction, *p* < 0.0001, *n* = 9).

**Figure 2:**
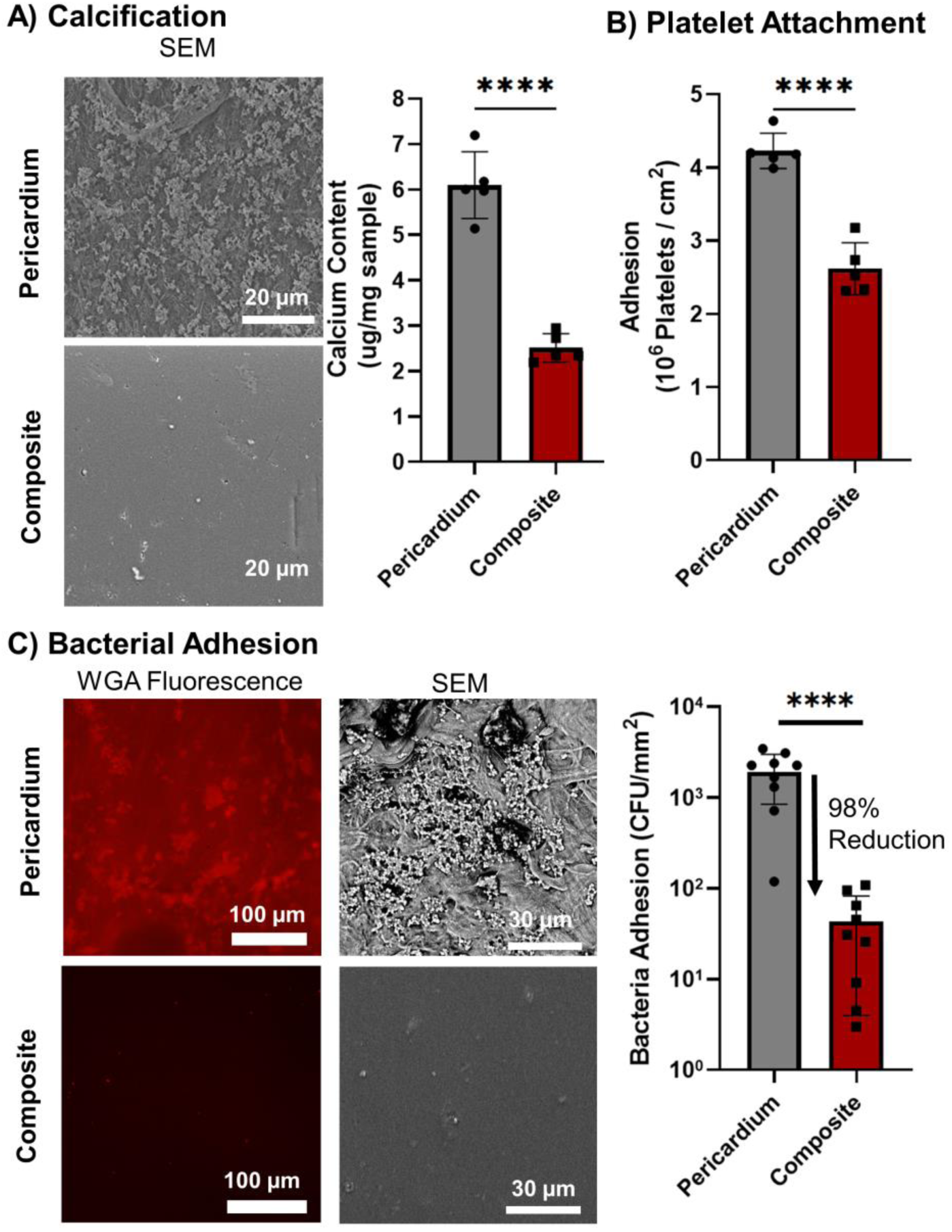
*In vitro* testing of biological responses of flat hydrogel-polyurethane fiber composites in comparison to glutaraldehyde-fixed pericardium control. A) Electron micrographs of the composite and pericardium specimens with quantification of calcium deposition during *in vitro* calcification (n = 5). Data is presented as the deposition of calcium in which the calcium content of non-treated samples was subtracted from the treated specimen calcium content to obtain the increase in calcium occurring from the exposure to the 2xSBF. B) Quantification of platelet attachment on the pericardium and composite samples (n = 5). C) Fluorescent and SEM micrographs of the bacterial adhesion to the pericardium and composite samples with quantification (n = 9). **** denotes statistical significance (p < 0.0001).

### 3.2 Heart Valve Fabrication and Assessment

For device-level assessment, electrospun tubes with a random fiber microarchitecture were fabricated with average thickness of 0.27 ± 0.02 mm and fiber diameter of 1.8 ± 0.3 µm. Suture retention strength of the heart valve material was 207 ± 8 gF. These values are greater than the suture retention strength reported by Choe et al. for decelluarized pericardium and native aortic leaflets.^44^ Our established redox coating system was used in combination with a custom 3D printed chamber and hanger to coat the electrospun tubes with a uniform IPN hydrogel with a thickness of 0.16 ± 0.02 mm, **Figure 3A**. Total composite thickness was 0.58 ± 0.02 mm. A custom expandable stent was utilized for valve assembly with the composite tube sutured to the custom stent to obtain uniform leaflets with full coaptation and minimal pinwheeling, **Figure 3B**.

**Figure 3:**
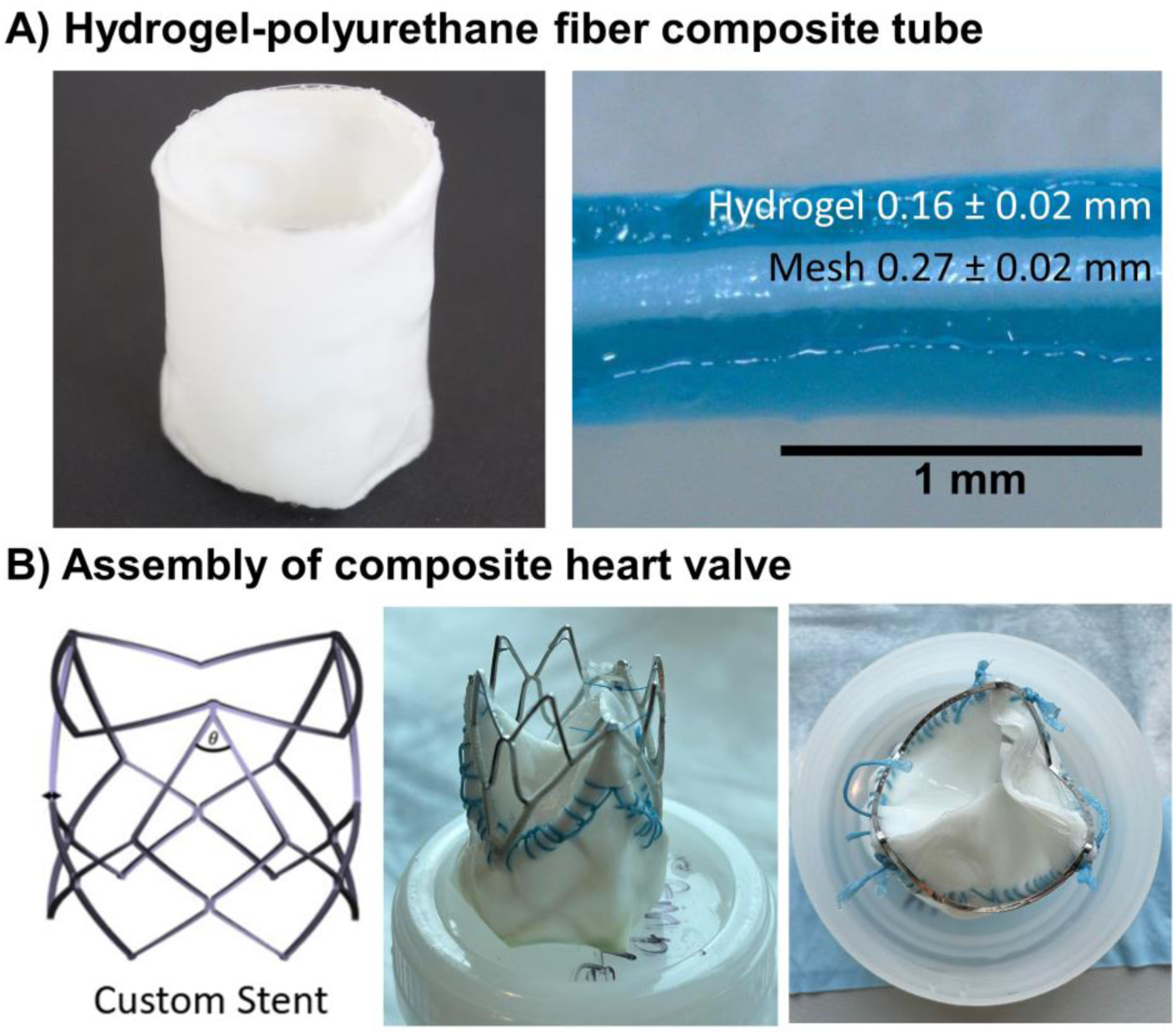
Assembly of the hydrogel-polyurethane fiber composite heart valve. A) Electrospun composite tube with cross-section showing the thicknesses of the mesh and hydrogel. B) Custom stent and assembled valve.

### 3.3 Hemodynamic Testing of Composite Heart Valves

The assembled valves underwent hemodynamic testing in a left heart simulator, **Figure 4** Valve performance was considered comparable and repeatable among the three valves assembled and tested. Each valve displayed opening and closing functions that mimics the behavior of native and prosthetic heart valves. The valves had an average effective orifice area (EOA) of 1.52 ± 0.34 cm^2^ and regurgitation fraction (RF) of 9.6 ± 1.8 %, which met requirements of ISO 5840-2:2021 standards for a 23 mm valve (EOA ≥ 1.25 cm^2^ and RF ≤ 10%).^56^ In addition, composite valves displayed a mean transvalvular pressure gradient comparable to current clinical valves (12.75 ± 4.39 mmHg).^57,58^

**Figure 4:**
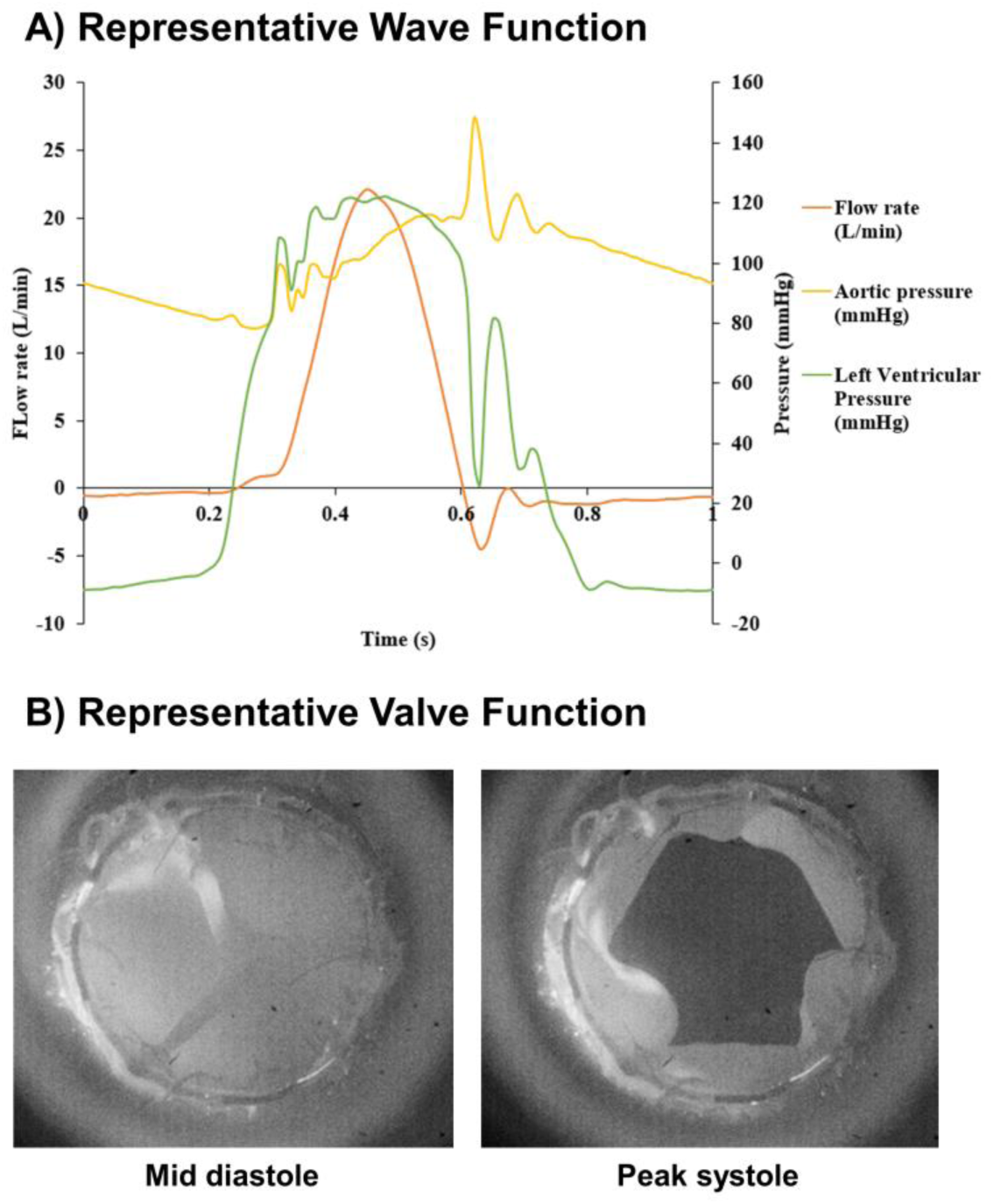
Hemodynamic assessment of the composite heart valve. A) Representative wave function during a single cycle for pressure and flow rate; B) Representative images of the valve fully closed and open during stages of a single cycle.

### 3.4 Control of Electrospun Microarchitectures

Rotational velocity was used to control fiber alignment, **Figure 5A**. Increasing the rotational speed from 50 RPM to 4000 RPM resulted in a marked increase in fiber alignment. A transition from randomly oriented fibers to highly aligned fibers with an associated increase in NOI from 14 ± 7 % to 85 ± 1 %, respectively. The NOI measured is well beyond physiological ranges (37-50%) demonstrating our ability to create highly aligned structures.^27,59^ Utilizing rotational velocity to impart alignment into electrospun mesh resulted in straight fibers in contrast to the tortuous fibers present in the native valve. To increase fiber tortuosity, heat-induced stress relaxation was utilized, **Figure 5B**. Removing the mesh from the mandrel during annealing (unconstrained), a wavy fiber orientation was achieved, whereas, annealing on the mandrel (constrained) retained the low tortuosity of the as-spun mesh. The tortuosity of the fibers for the unconstrained mesh was increased to 1.08 ± 0.04 as compared to the constrained mesh at 1.02 ± 0.01. This tortuosity is within the lower reported physiological ranges for heart valves.^60,61^ To introduce curvilinearity into the electrospun leaflets, the mandrel geometry was altered from a cylinder to a cone, **Figure 5C**. Utilizing a cone with a base diameter of 7.5 cm and a height of 3.5 cm, a change in the preferred angle of alignment across the material was measured at 1.9 ± 0.4 °/mm, compared to the cylinder at 0.3 ± 0.2 °/mm. This degree of curvilinearity approximates recorded literature heart valve curvilinear values.^27^

**Figure 5:**
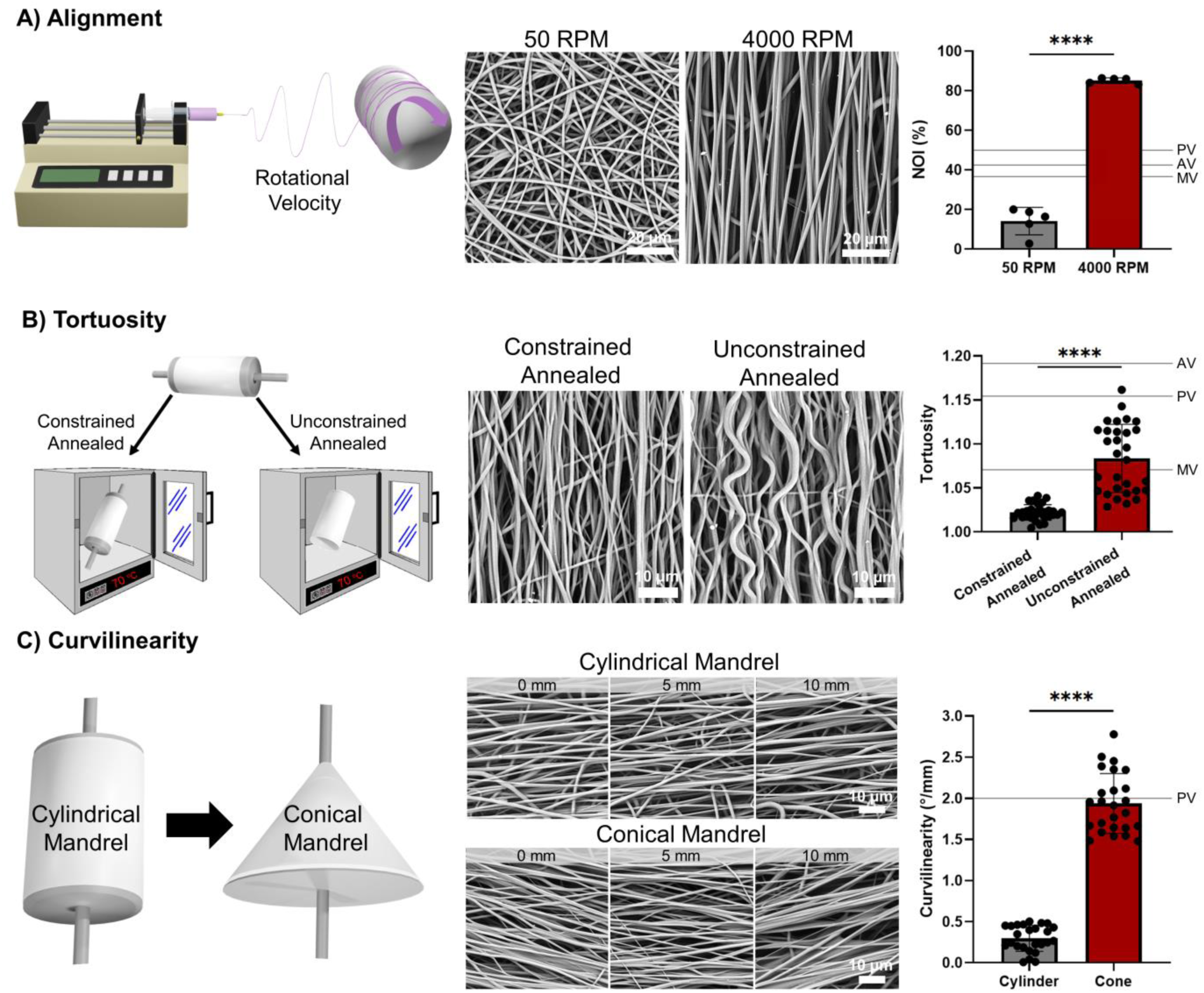
Microarchitectural control of electrospun polyurethane meshes. A) Control of alignment with rotational velocity (n = 5). Pulmonary valve (PV) literature value^27^, aortic valve (AV) literature value^27^, and mitral valve (MV) literature value^59^. B) Introduction of fiber tortuosity with post-fabrication annealing. Tortuosity was calculated according to Eq. 1, 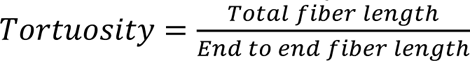 (n = 30). PV literature value^61^, AV literature value^61^, MV literature value^60^. C) Introduction of curvilinearity through changes in the mandrel geometry from a cylindrical geometry to a conical geometry. Micrographs were taken at 0 mm, 5 mm, and 10 mm along the section following the fiber alignment direction to illustrate how the main angle of alignment changes across the mesh. Curvilinearity was calculated according to Eq. 2, 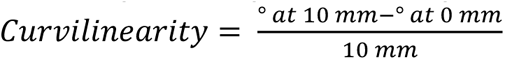 (n = 25). C) PV literature value^27^. **** indicates statistical differences, (p < 0.0000001).

## 4. Discussion

Despite concerted efforts over several decades, polymeric heart valve materials have yet to fully meet the complex demands of the cardiovascular environment. Designing a single-phase material for this space that is able to improve upon the durability of current bioprosthetic valves while retaining comparable hemocompatibility has remained elusive. Herein, we demonstrated the potential of a composite heart valve material composed of a poly(carbonate urethane) electrospun mesh that provides bulk mechanical properties and a tough hydrogel coating to confer antifouling properties.

The composite material was evaluated for its biological responses including platelet attachment, bacterial adhesion, and calcification propensity. All three biological responses were markedly improved as compared to glutaraldehyde-fixed pericardium demonstrating biological suitability for valve applications. Platelet attachment on the composite was 2.6 x 10^6^ platelets/cm^2^ compared to the pericardium at 4.3 x 10^6^ platelets/cm^2^. Current bioprosthetic valves do not require patients to be on long-term anticoagulant therapy and as such, this 38% reduction in platelet attachment compared to the glutaraldehyde-fixed control is anticipated to confer an even greater degree of thromboresistance. This finding is even more significant in light of investigations into subclinical thrombus in tissue based bioprosthetic valves that are suggested to have a role in major failures mechanisms such as structural degeneration.^62,63^ Significantly, areas of calcification in not only bioprosthetic valves, but also in attempts in previous polymeric valves have been correlated with subclinical microthrombus that may provide initial nucleation sites.^64^ Thus, the enhanced thromboresistance serves not only to mitigate thrombus based failure, but also aid in mitigating calcification-based failure. In addition to the hydrogel coating conferring thromboresistance, there was a marked resistance to bacterial adhesion with a 98% reduction in colony forming units as compared to the glutaraldehyde-fixed pericardium control. This degree of bacterial resistance is significant given that bacterial endocarditis results in valve failure in up to 2.3% of patients per year, with a majority being staph based infecitons.^65–67^ Although these *in vitro* assessments provide preliminary demonstration of the antifouling nature of the hydrogel coating, it is necessary to continue evaluations under more rigorous assessments such as flow loop studies and long-term *in vivo* assessments due to the complex nature of thrombus generation and fouling mechanisms that are not replicated in static assessments.

In addition to biofouling-based biological failures, material calcification is known to accelerate structural degeneration and cause leaflet embrittlement.^68^ The calcification propensity of the composite material was tested using a concentrated simulated body fluid. At the end of the two-week study, the composite displayed a 59% decrease in calcium deposition as compared to the glutaraldehyde-fixed pericardium. Scanning electron microscopy revealed significant calcium nucleation on the pericardial samples, whereas, the composite remained similar to the untreated control, **Figure 3A** and **Figure S1**. This *in vitro* assessment demonstrates an innate reduction in material calcification propensity for the composite compared to the pericardium that may lend to an improved long-term durability. However, investigation with dynamic studies and *in vivo* assessments are warranted to further evaluate the material. Dynamic assessments have shown larger amounts of calcium deposition compared to static assessments.^69^ Similarly, *in vivo* calcification is much more complex than simple innate material chemistry driving deposition as biological responses such as protein adsorption and platelet attachment could serve as nucleation sites for calcification as discussed prior.^64^ Overall, these initial assessments indicate biological suitability of the hydrogel-polyurethane fiber composites.

Although flat sheets permitted the assessment of *in vitro* biological responses, full valve constructs were needed for hemodynamic testing. We previously developed a redox-based coating method to overcome the limitations of molds during hydrogel coating, including limited substrate geometries and uneven coatings due to poor positioning within the mold.^23,33,70^ Due to its promise in providing uniform and thin coatings, this redox method was utilized to fabricate the composite heart valves. First attempts resulted in uneven coatings due to buckling of the heart valve tubes during coating and contact with the container wall. To improve the uniform coating of large tube constructs, a custom 3D printed chamber and hanger system was developed. The chamber with a central pillar allowed for a reduction in the polymer used. The hangers stabilized the mesh to prevent buckling during submersion for coating and guided the mesh within the chamber to mitigate wall contact. Utilizing this approach, a hydrogel thickness of 0.16 ± 0.02 mm was achieved demonstrating the ability to create thin uniform coatings on large constructs. The thin coating resulted in a total composite thickness of 0.58 ± 0.02 mm, which is comparable to current prosthetic valves such as the Carpentier-Edwards PERIMOUNT Magna that has a leaflet thickness of 0.56 mm.^71^ Overall, the 3D printed chamber and hanger systems in combination with our redox coating system permitted the fabrication of uniform hydrogel coatings on large and complex substrate geometries thereby enabling hemodynamic assessments.

Heart valves were assembled from the hydrogel-polyurethane fiber composite tubes by suturing the tube to a custom cobalt chromium stent. Hemodynamic performance was assessed in a left heart simulator. As a first assessment, the performance of the composite valve met requirements of ISO 5840-2:2021 standard for 23 mm diameter valves (EOA ≥ 1.25 cm^2^ and RF ≤ 10%)^56^ and the EOA was comparable to current clinical valves.^58,72^ An adequate EOA is necessary to minimize the pressure gradients across the valve that can negatively reduce patient outcomes and lead to events such as congestive heart failure.^73,74^ In addition, the mean transvalvular pressure gradient (12.75 ± 4.39 mmHg) was comparable to current clinical valves such as the Medtronic Hancock (16.48 mmHg), Medtronic Mosaic (14.64 mmHg), and St. Jude Epic (13.48 mmHg).^58^ Overall, the hemodynamic performance was similar to clinical valves and meets currents ISO standards, which support the use of the hydrogel-polyurethane fiber composite as a synthetic heart valve replacement option.

Although initial benchtop testing is promising, accelerated wear testing and large animal testing with appropriate size and host response are needed to assess the long-term performance of the valve. We anticipate that the random microarchitecture used here as proof-of-concept is likely to possess regions of stress concentration that may impact long-term durability compared to bioinspired microarchitectures. Indeed, isotropic heart valves have been shown to have higher principal stress compared to anisotropic leaflets.^75–77^ In addition, biomimetic fiber geometries have demonstrated improvements in strain homogeneity compared to simple fiber architectures.^28,29^ Combined, these studies indicate that bioinspired fiber microarchitectures may offer an additional design space to improve heart valve durability. To this end, we developed methodologies to precisely control electrospun fiber microarchitecture to expand the design space toward enhancing durability. The electrospinning methodologies developed for these studies demonstrated improved control of fiber microarchitectures of alignment, curvilinearity, and tortuosity that meet or exceed native valve ranges. The broad range of fiber alignment achieved greatly surpassed the degree of alignment for the native valve that is recorded between ∼35-50%.^27,59^ We have previously shown similar degrees of alignment produce a high degree of mechanical anisotropy.^21^ Intermediate rotational velocities can be used to reduced degrees of alignment and control the degree of anisotropy as needed. Tortuosity was imparted through post-fabrication annealing treatments. Electrospinning induces internal stresses within the fiber.^19^ Annealing provides energy to promote polymeric chain mobility toward releasing the internal stress. Annealing the mesh while it is on the mandrel, in a constrained manner, the internal stress is released without permitting fiber contraction along the length thereby maintaining the straight fiber presentation. By removing the mandrel and then annealing, fibers contract along their length and buckle producing a crimped architecture. Utilizing this approach, we were able to obtain an increase in tortuosity from 1.02 ± 0.01 to 1.08 ± 0.04 that is within the lower bounds of reported valves fiber crimp. Fiber crimp has been shown to delay strain hardening responses that can be used to induce a non-linear stress-strain response similar to the native heart valves.^78^ Lastly, curvilinearity was imparted through mandrel geometry changes from a cylinder to a cone. The pitch of the cone was utilized to obtain a change in the main angle of alignment, 1.9 ± 0.4 °/mm, that is consistent with the pulmonary valve based on microarchitecture assessments by Joyce et al.^27^ As Hobson et al. demonstrated, curvilinearity can be utilized to improve strain homogeneity that is hypothesized to improve long-term durability by reducing stress concentrations that can contribute to material fatigue.^28^ Alterations in cone pitch can be designed to adjust the change in main angle of alignment with increased pitches increasing the change in the main angle of alignment across the length of the leaflet. We hypothesize that a synthetic heart valve that possesses aspects of these bioinspired microarchitectures will improve the durability compared to a random fiber microarchitecture by recapturing critical aspects of the native valve mechanical properties.

Although prior research supports that these bioinspired microarchitectures can enhance valve mechanical properties to confer enhanced durability, the multidimensional fiber parameter space creates a large number of design combinations. Iterative testing to identify target graft parameters with optimal properties would be time-consuming even with the previously identified structure-property relationships. In contrast, coupled computational mechanics and numerical optimization strategies provide efficient approaches to iterate through multi-dimensional parameter space *in silico* and identify favorable parameter combinations to accelerate device design. The advanced fabrication techniques developed here can then be utilized to fabricate valves with target parameters and enhanced durability.

## 5. Conclusion

In this work, we evaluated a composite material composed of an electrospun polyurethane mesh and hydrogel coating as a tri-leaflet replacement heart valve. The hydrogel coating of the polyurethane mesh was used to impart favorable biological responses to the candidate valve material. Platelet and bacterial attachment were reduced compared to the pericardium demonstrating the antifouling nature of the hydrogel coating. There was also a notable reduction in the calcification of the composite material. A custom 3D printed coating setup was used to make valve composites for hemodynamic testing. Regurgitation fraction and effective orifice area met ISO 5840-2:2021 requirements and the mean pressure gradient was comparable to current clinical valves. This combination of biological suitability and hemodynamic performance support the feasibility of this hydrogel-polyurethane fiber composite as a synthetic heart valve material. However, it is anticipated that the isotropic mechanical properties produced by the random microarchitecture will possess high peak stresses that may impact long-term durability. To this end, advanced electrospinning methodology was used to attain polyurethane fiber microarchitecture that achieves or exceeds the fiber alignment, curvilinearity, and tortuosity ranges present in the native valve that is anticipated to improve strain homogeneity and reduce peak stresses for enhanced durability. Future work will utilize model-directed design in combination with this advanced electrospinning methodology to optimize durability as a function of fiber microarchitecture.

## Acknowledgment

This work was supported by the American Heart Institute Predoctoral Fellowship (23PRE1020036 – DOI: https://doi.org/10.58275/AHA.23PRE1020036.pc.gr.161179), NSF GRFP (2023355666), and NSF GRFP (2020300397). Bionate® 80A was provided by DSM Biomedical (Berkeley, CA).

## Data Availability Statement

All data is available on request from the authors.

---

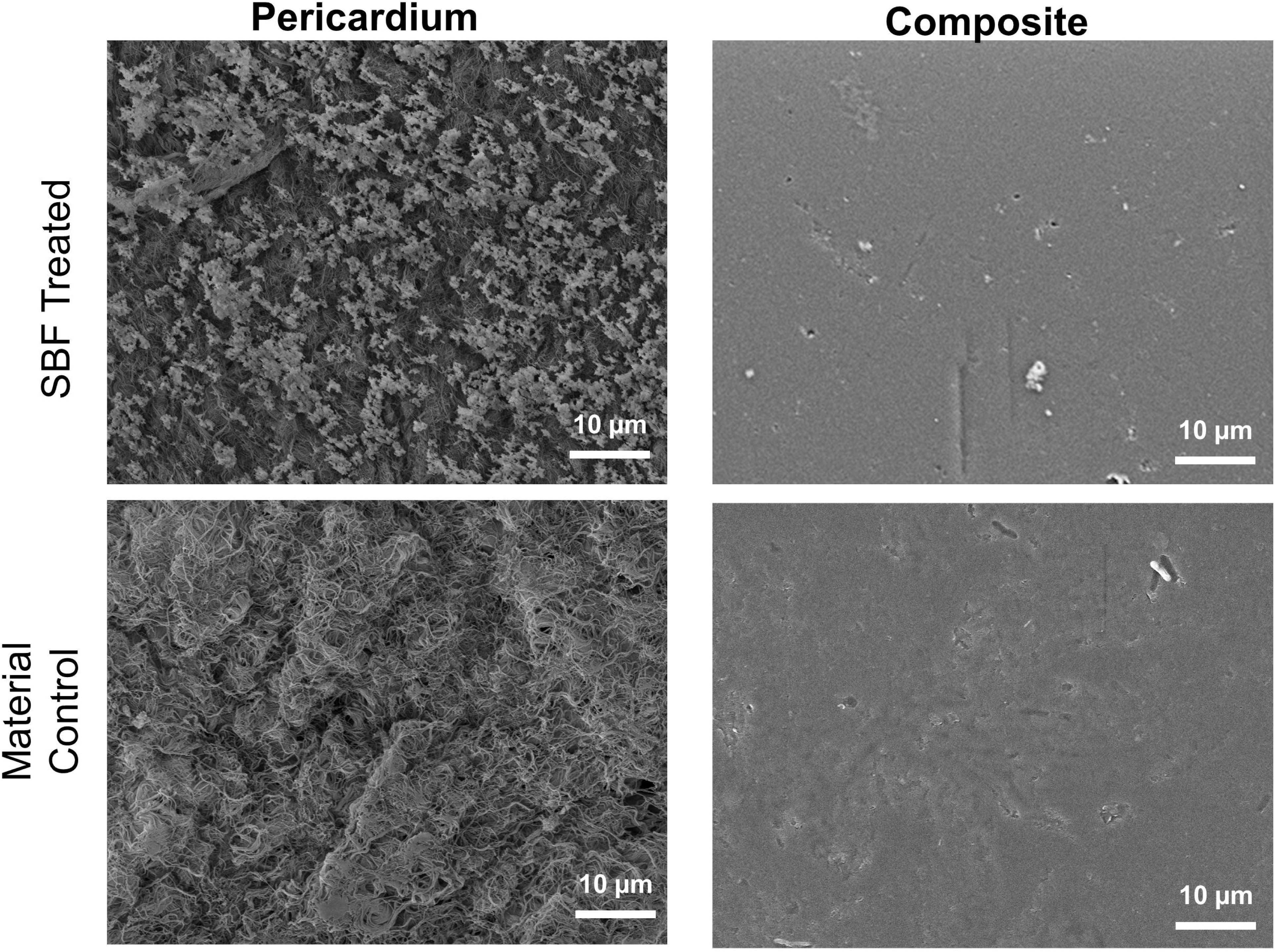

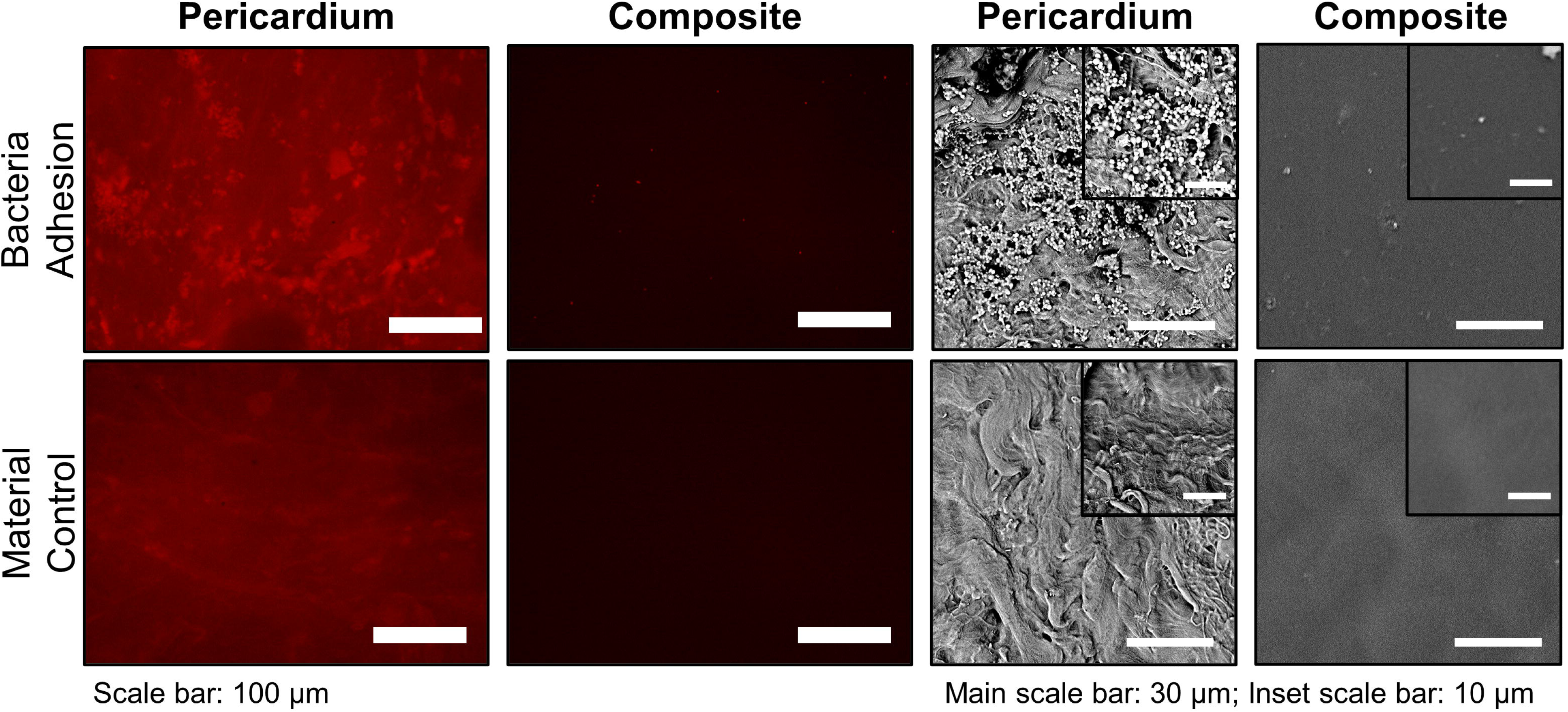

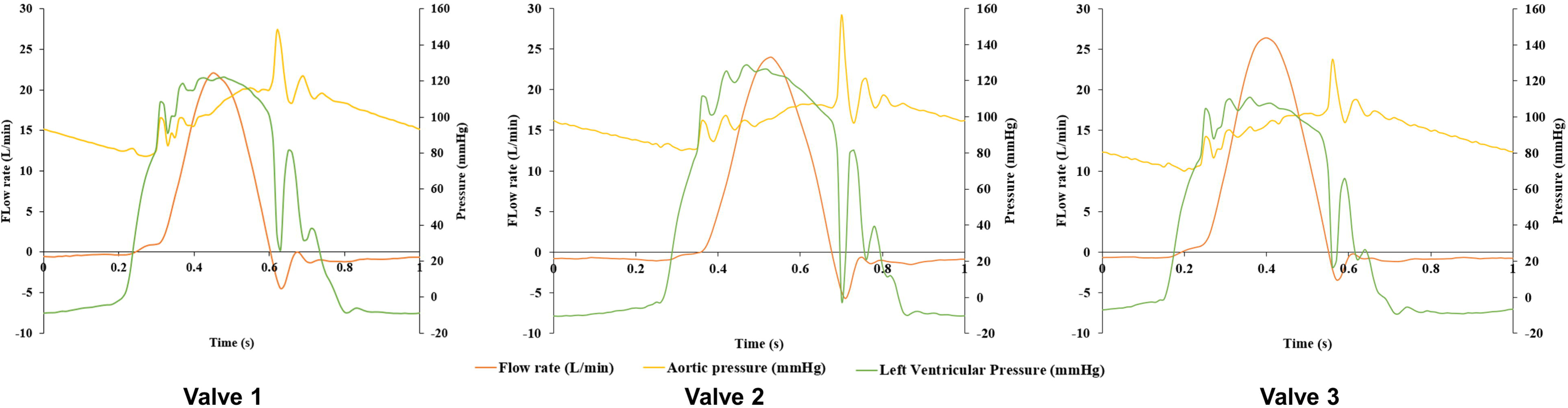

