## Supplemental Files for "Hydrogel-polyurethane fiber composites with enhanced microarchitectural control for heart valve replacement"

**Supplemental Figures**

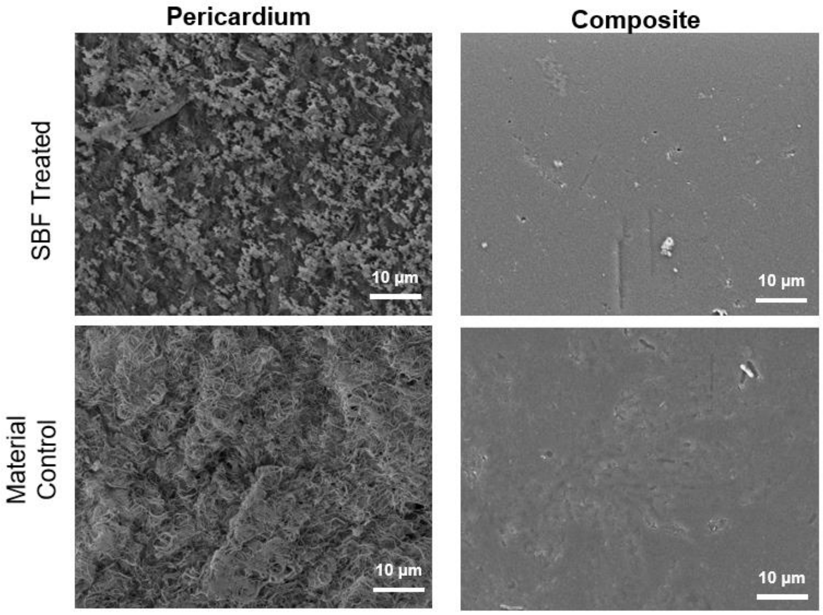


**Figure S1**: Simulated body fluid treated samples for calcification propensity of the pericardium and composite verse their material controls.

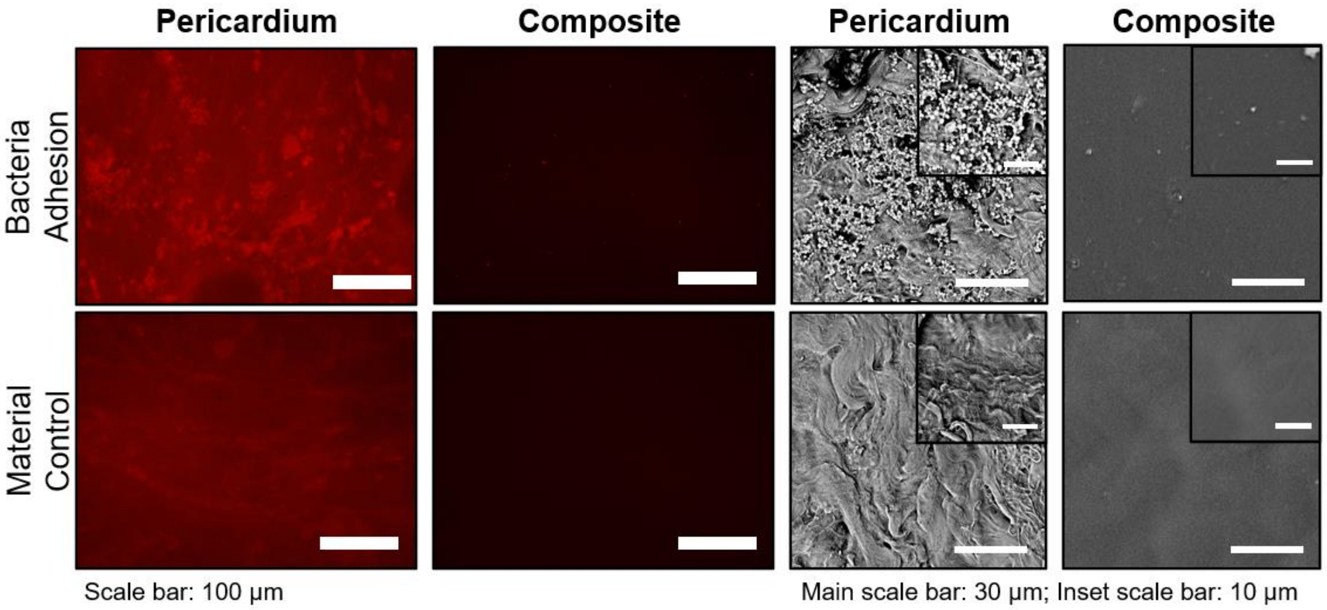


**Figure S2**: Bacterial adhesion for the pericardium and composite verse their material controls.

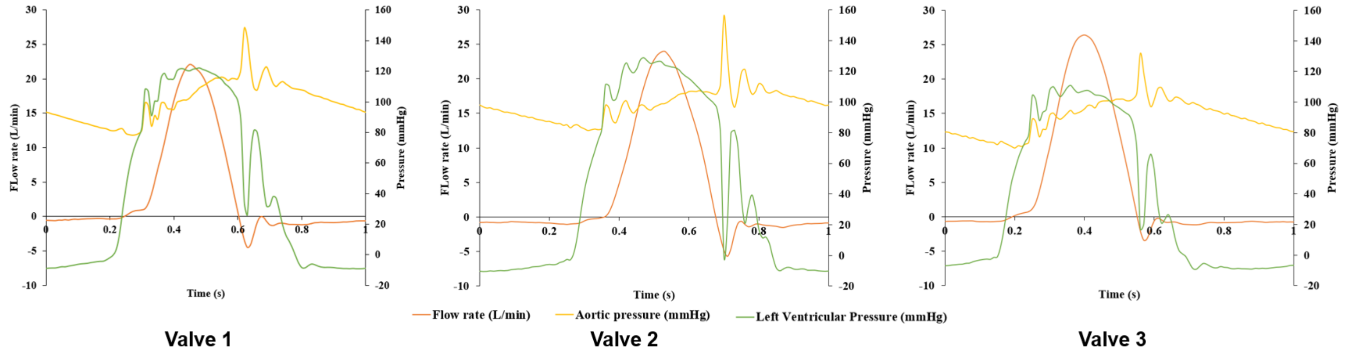


**Figure S3**: Representative wave functions from each of the three assembled hydrogel-polyurethane fiber composite heart valves.
